# Targeting Galectin-3 to modulate inflammation in LAMA2-deficient congenital muscular dystrophy

**DOI:** 10.1101/2025.03.12.642905

**Authors:** Yonne Karoline Tenorio de Menezes, Jinseo Lee, Jia Qi Cheng-Zhang, Marie A. Johnson, Ruvindi N. Ranatunga, Dwi U. Kemaladewi

## Abstract

LAMA2-deficient congenital muscular dystrophy (LAMA2-CMD) is a severe neuromuscular disorder characterized by muscle degeneration, chronic inflammation, and fibrosis. While inflammation is one the hallmarks of LAMA2-CMD, the immune cell composition in laminin-deficient muscles remains understudied. Consequently, targeted pharmacological intervention to reduce inflammation remains underexplored.

Here, we characterized the immune landscape in the dyW mouse model of LAMA2-CMD using RNA sequencing and flow cytometry. Transcriptomic analysis of dyW quadriceps femoris muscle identified 2,143 differentially expressed genes, with most upregulated genes linked to immune-related pathways. *Lgals3* (Galectin-3) was significantly upregulated and identified as a key upstream regulator of the immune-related pathways. Flow cytometry revealed elevated leukocyte (CD45⁺) infiltration, with macrophages as the predominant population. Pro-inflammatory (M1) macrophages were increased, whereas anti-inflammatory (M2) macrophages remained low, indicating persistent and unresolved inflammation. Notably, Galectin-3^+^ macrophages were significantly enriched, suggesting that Galectin-3 drives inflammation in LAMA2-CMD.

Treatment of dyW mice with TD-139, a Galectin-3 inhibitor, reduced leukocyte infiltration, decreased Galectin-3^+^ macrophages, and shifted macrophage polarization toward an M2 anti-inflammatory profile. RNA sequencing of TD-139-treated dyW muscles showed upregulation of muscle contraction pathways and downregulation of fibrosis-related genes. These findings highlight Galectin-3^+^ macrophages as key contributors to LAMA2-CMD pathophysiology and support further exploration of TD-139 as a potential therapeutic strategy for LAMA2-CMD and other dystrophic conditions driven by chronic inflammation.

## Introduction

Congenital muscular dystrophy encompasses hereditary diseases that primarily affect skeletal muscle and become apparent at or near birth. Within this group, congenital muscular dystrophy type 1A (LAMA2-CMD; MIM: 607855) emerged as one of the most prevalent and severe, with no treatment identified to date (1). LAMA2-CMD is caused by mutations in the *LAMA2* gene encoding the laminin α2 (merosin) subunit of the laminin-211 heterotrimeric protein, which plays a crucial role in providing stability to muscle fibers and maintaining their homeostasis (2–5). Individuals with LAMA2-CMD experience muscle weakness and hypotonia from birth (1). As the disease progresses, severe muscle wasting, joint contractures, significant impairment in motor function, and respiratory difficulties profoundly impact the life expectancy and well-being of young patients (2–6). The deficiency in laminin α2, either complete or partial, results in muscle fiber atrophy, necrosis and apoptosis of muscle cells, extensive infiltration of connective tissue, and early inflammation (6–8).

Inflammation plays a dual role in muscular dystrophies, including LAMA2-CMD and Duchenne muscular dystrophy (DMD)—initially facilitating muscle repair, but becoming detrimental when chronic, driving fibrosis and disease progression (7, 9–13). In LAMA2-CMD, unlike other muscular dystrophies, the inflammatory infiltrates peak early and decline during the later stages (7, 13). A better understanding of the inflammatory dynamics may inform development of new therapeutic strategies for treating LAMA2-CMD.

Pegoraro *et al* conducted the first histopathological analysis in muscle biopsies from LAMA2-CMD patients and identified the presence of inflammatory cells around blood vessels and muscle fibers (7). Notably, they also observed that inflammation detected in individuals at the age of 5 months was absent in follow-up biopsies taken 9 months later, suggesting a decline in inflammation with age. This finding was largely recapitulated in a subsequent study by Konkay *et al*, in which muscle biopsies from 65 patients aged between 10 days and 11 years showed variable degrees of inflammation (13). These studies provide important clinical evidence of a neonatal inflammatory processes associated with LAMA2-CMD; however, due to the scarcity of patient materials, the composition of resident immune cells was not quantitatively analyzed.

In contrast to the rarity of human muscle biopsies, various mouse models to study LAMA2-CMD are available, therefore becoming invaluable tools to study disease pathogenesis. From mouse models, it has been reported that no signs of apoptosis, inflammation, or excessive extracellular matrix deposition are observed during embryonic development, indicating that LAMA2-CMD muscle pathology emerges postnatally (8, 14). Histopathological changes appear as early as postnatal day 1, with mononuclear cell infiltration detectable between 4 days and 1 week after birth (8). Among the infiltrating cells, macrophages have been identified as one of the predominant immune cell populations infiltrating the affected muscle tissues in LAMA2-CMD across different stages of the disease in mouse models with different levels of LAMA2 expression (8, 15–18).

Macrophages are essential for muscle repair and regeneration, orchestrating inflammation, clearing damaged cells through phagocytosis, and facilitating tissue remodeling (19, 20). They act as key regulators of the healing process through a mechanism known as polarization, where they adopt different functional states in response to environmental signals. These states are broadly classified into M1 (pro-inflammatory) and M2 (anti-inflammatory), each playing distinct roles in immune responses, tissue repair and fibrosis. Persistent M1 activation exacerbates muscle damage by producing pro-inflammatory cytokines, while M2 macrophages support tissue repair and regeneration. The balance between M1-driven inflammation and M2-mediated repair is crucial in muscular dystrophy, and dysregulation can lead to excessive inflammation and fibrosis instead of effective regeneration (9, 10, 20–25). However, the dynamics of macrophage polarization in LAMA2-CMD remains unknown.

In this study, we conducted a comprehensive characterization of the immune cell infiltrate in LAMA2-CMD muscles using RNA sequencing and flow cytometry-based immunophenotyping. Transcriptomic analysis identified *Lgals3* (Galectin-3) as a key upstream regulator, strongly linked to inflammatory and fibrotic pathways. Flow cytometry further revealed a transient yet robust immune response, with peak leukocyte infiltration at 2 weeks, predominantly driven by macrophages.

Given the high abundance of Galectin-3^+^ macrophages and their association with fibrotic pathways, we treated LAMA2-CMD mice with TD-139, a small-molecule Galectin-3 inhibitor currently under clinical investigation for fibrotic and inflammatory diseases (26–30). TD-139 treatment significantly reduced leukocyte infiltration, altered macrophage polarization, and shifted transcriptomic profiles toward pathways associated with muscle contraction and adaptation, while simultaneously downregulating fibrosis-associated genes.

This study represents the first systematic integration of transcriptomic analysis and flow cytometry-based immunophenotyping to characterize the immune landscape of LAMA2-CMD. Our findings position Galectin-3 as a critical modulator of macrophage-driven inflammation and fibrosis, providing a strong rationale for its therapeutic targeting. These results lay the foundation for future studies to optimize treatment strategies and evaluate long-term therapeutic outcomes, not only in LAMA2-CMD but also in other dystrophic conditions where chronic inflammation is a key driver of disease progression.

## Results

### Transcriptomic profiling reveals muscle dysfunction and immune activation in dyW mice

First, we analyzed the transcriptomic profiles of quadriceps femoris muscles isolated from 5-week-old wildtype (WT) and LAMA2-deficient dyW mice, a genetic mouse model of LAMA2-CMD (**Fig. 1A**). A total of 2,143 differentially expressed genes (DEGs) were significantly altered between the dyW and WT mice. Of these, 1,409 genes (65%) were upregulated (Log_2_ fold-change ≥1.5 and a false discovery rate (FDR) ≤0.05), and 734 (35%) were downregulated (Log_2_ fold-change ≤ −1.5 and a false discovery rate (FDR) ≤0.05) (**Fig. 1A**; **Supplemental Table 1**). Unsupervised hierarchical clustering demonstrated a clear separation in gene expression profiles between the dyW muscle and WT control (**Fig. 1B**), indicating that *Lama2* deficiency has apparent consequences on the quadriceps transcriptome.

**Figure 1:**
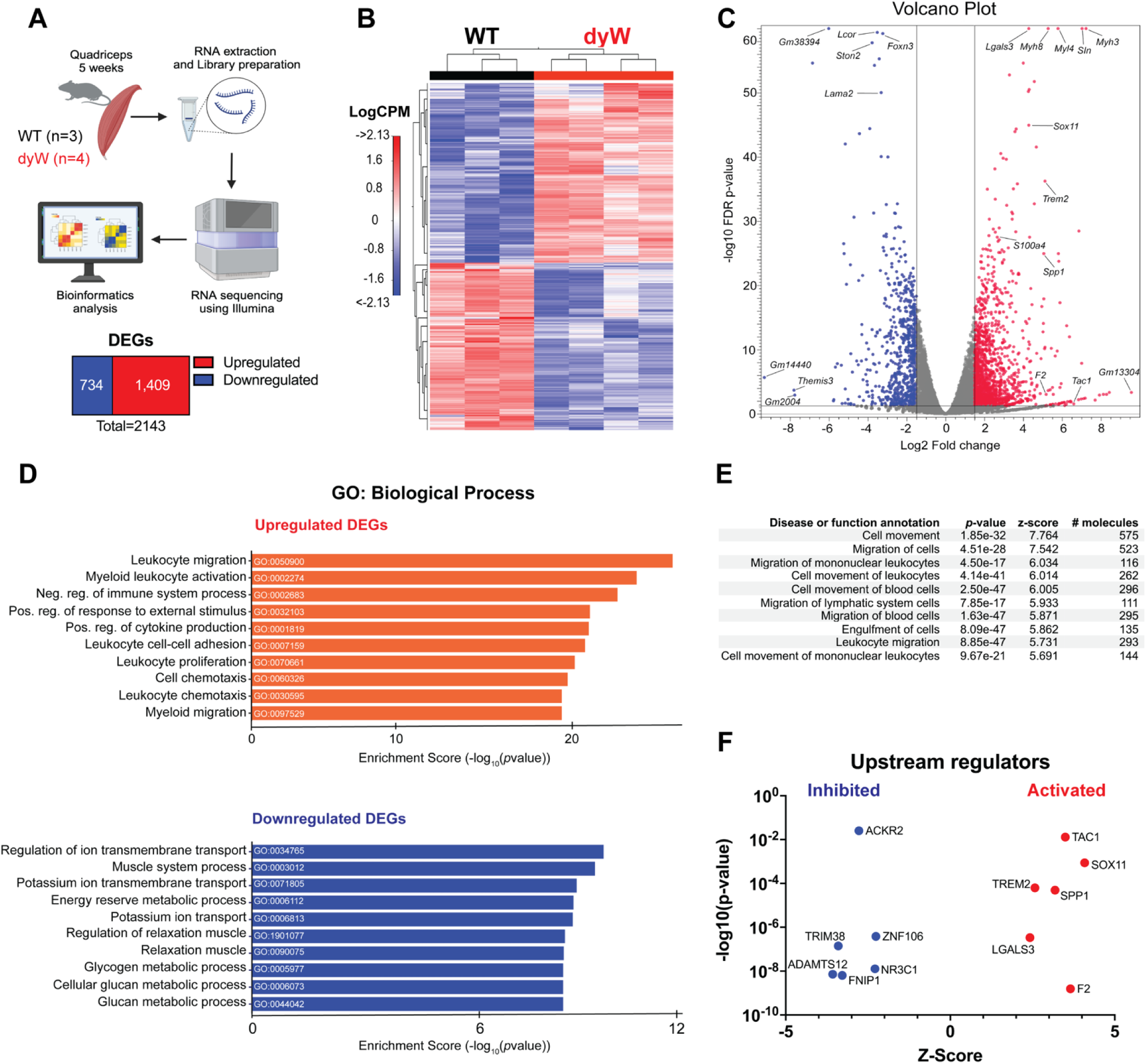
Transcriptomic analysis of quadriceps from dyW and wildtype (WT) at 5 weeks of age. (A) Experimental workflow. Quadriceps were harvested from 5-week-old WT and dyW mice (WT n=3, dyW n=4). RNA was extracted and libraries were prepared for RNA sequencing using Illumina technology. The sequencing data were subjected to bioinformatic analysis to identify differentially expressed genes (DEGs). (B) Heatmap shows hierarchical clustering of differentially expressed genes (DEG) between WT and dyW mice. Rows represent individual genes, and columns represent individual samples. The LogCPM colors indicate expression levels across samples (red = high, blue = low). (C) Volcano plot displays DEGs in dyW versus WT with the most significant DEGs highlighted. Red dots represent up-regulated genes and blue dots represent down-regulated genes, and the greys dots represents non-significant genes. The x-axis shows the log_2_ fold change, and the y-axis shows the −log10 FDR p-value. (D) Gene ontology (GO) analysis with a focus on biological processes displays the top 10 significantly enriched upregulated (orange) and downregulated (blue) processes in DEGs. (E) Disease and function annotations from Qiagen’s Ingenuity Pathway Analysis (IPA) showing the top 10 significant disease or function annotations among the DEGs. (F) Upstream regulator analysis using IPA software. Top 6 most significantly activated and inhibited regulators ranked by the magnitude of activation z-score. An activation z-score of >2 indicates predicted to be activated, and less than −2 is predicted to be inhibited.

The volcano plot in **Fig. 1C** shows the most statistically significant DEGs in dyW vs. WT quadriceps. Notably, developmental isoforms of myosin heavy chain (Myh) and myosin light chain (Myl), such as *Myh8*, *Myh3*, and *Myl4,* were significantly upregulated. Additionally, inflammatory genes such as *Lgals3, Trem2* and *Spp1* were also significantly upregulated. *Lgals3* and *Spp1* have been previously associated with dystrophic muscle conditions, including Duchenne and LAMA2-CMD, where they contribute to pro-inflammatory and pro-fibrotic responses (18, 31, 32).

To pinpoint the biological pathways linked to pathogenic mechanisms in dyW muscles, we used SRplot database (33) to conduct conducted Gene Ontology (GO) analysis on the upregulated and downregulated DEGs with a focus on biological processes (**Fig. 1D**, **Supplemental table 2).** The GO analysis demonstrated that the upregulated DEGs were enriched in leukocyte migration, myeloid activation, regulation of immune system process, proliferation, and cell chemotaxis, all of which are immune-related terms. In contrast, the downregulated DEGs were enriched in the regulation of ion transmembrane transport, muscle system process, regulation of relaxation of muscle, energy reserve, and glucan metabolism, suggesting an impairment in overall muscle function.

To further explore these findings, we analyzed the DEGs using Ingenuity Pathway Analysis (IPA) focused on biological ontology, such as disease and functions, to identify which functional categories are most strongly impacted by *Lama2* mutation. The top three functions by activation z-score identified by this analysis were cell movement, migration of cells, and migration of mononuclear leukocytes (**Fig. 1E**). Subsequently, we performed upstream regulator analysis, which predicts key molecules that regulate gene expression changes in our transcriptomic data, using IPA. This analysis identified of 68 regulators that all showed a direction of activity consistent with the activation state (**Supplemental Table 3**). The six regulators with the highest activation scores included both activated and inhibited regulators. The activated regulators were *Sox11*, *F2*, *Tac1*, *Spp1*, *Trem2*, and *Lgals3*, while the inhibited regulators were *Adamts12*, *Trim38*, *Fnp1*, *Ackr2*, *Nr3c1*, and *Znf106* **(Fig. 1F**).

Taken together, these findings highlight a strong inflammatory response driven by *Lama2* mutation in LAMA2-CMD, linking immune cell infiltration and activation to muscle dysfunction and identifying key inflammatory mediators as potential therapeutic targets.

### Skeletal muscles from dyW mice exhibit increased leukocyte (CD45^+^) infiltration throughout disease progression

Given the enrichment of pathways related to leukocyte migration and myeloid activation in dyW muscles, we sought to investigate whether immune cell infiltration follows a specific spatiotemporal pattern. To explore the dynamics and composition of these immune cells, we performed flow cytometry analysis on dyW and WT mice hindlimbs at different time points between 2 and 5 weeks of age (**Fig. 2A**). The leukocyte count expressed as frequency, assessed using the pan-leukocyte marker CD45, was significantly higher in dyW muscles compared to age-matched WT controls, with peak of infiltration observed between 2 and 4 weeks. In contrast, CD45^+^ cell frequencies remained consistently low in WT mice (**Fig. 2B**).

**Figure 2:**
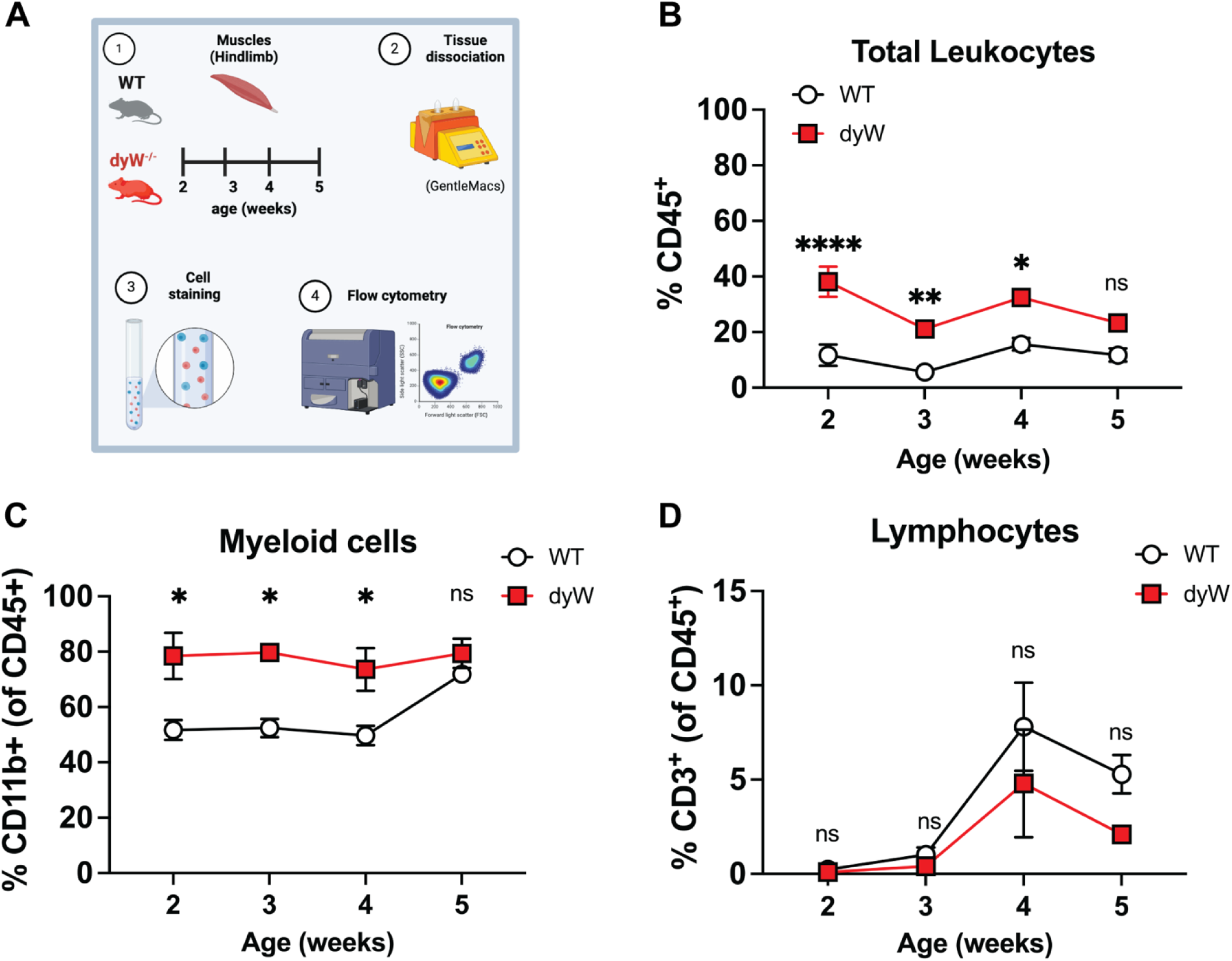
Dynamics of Leukocyte (CD45^+^) Infiltration in dyW Muscles. (A) Schematic representation of the study design and methodology used to immunophenotype hindlimbs from wildtype (WT) and dyW mice. (B) Frequencies of CD45^+^ cells in hindlimbs from WT and dyW mice at the indicated ages. (C) Frequencies of CD11b^+^ and (D) CD3^+^ cells as a percent of CD45^+^ cells at the indicated ages. n= 5-7 mice/group. Data are presented as mean ± SEM. Results are pooled from 3 or more independent experiments. Statistical analysis was determined using two-way ANOVA with Tukey’s multiple comparisons. ns=not significant, *p ≤ 0.05, **p ≤ 0.01, ****p ≤ 0.0001.

The GO analysis also revealed a significant enrichment in myeloid migration pathways. Consistent with this, further analysis revealed an increased frequency of myeloid cells (CD11b^+^) in dyW muscles compared to WT (**Fig. 2C**). In contrast, while lymphocytes (CD3^+^) were detected in both dyW and WT muscles, their frequencies remained consistent across groups, showing no statistically significant differences (**Fig. 2D**). Given the presence of lymphocytes in both dyW and WT muscles, we further analyzed T cell subsets to determine whether specific populations were altered in the disease context (**Supplemental Fig. 2)**. While CD4⁺ and CD8⁺ T cell counts showed no statistically significant differences between groups, remaining at levels similar to those observed in WT-controls, we observed an early increase in activated regulatory T cells (Tregs) in dyW muscles, peaking at 2 to 3 weeks before declining. These Tregs displayed an activated phenotype, with higher frequencies of GITR^+^KLRG1^+^ Tregs in dyW muscles compared to WT (**Supplemental Fig. 2)**. This suggests that Tregs may play a role in modulating the early inflammatory response in LAMA2-CMD.

Taken together, these data suggest that dyW muscles exhibit a dynamic immune response, marked by fluctuating CD45⁺ cell frequencies, a predominant myeloid presence, and an early increase in activated Tregs. The persistent infiltration of these immune cells, likely driven by ongoing muscle injury and degeneration, suggests a sustained inflammatory response that might contribute to disease pathology. These findings highlight myeloid cells as key mediators of inflammation and potential targets for therapeutic intervention.

### Macrophages in dyW muscles are increased and express high levels of Galectin-3

Myeloid cells were the most abundant immune population in dyW muscles; therefore, we sought to investigate their role in disease pathology, with a focus on macrophages due to their crucial role in muscle repair and homeostasis (34, 35). While macrophages are known to orchestrate both inflammatory and regenerative responses following acute muscle injury (35), their role in LAMA2-CMD remains understudied. We hypothesize that macrophage balance is disrupted in LAMA2-CMD, favoring a pro-inflammatory (M1) state that sustains chronic inflammation rather than an anti-inflammatory (M2) state that supports muscle regeneration (9, 24). To test this hypothesis, we quantified the total macrophage population and analyzed the frequencies of M1 and M2 macrophages in WT and dyW mice from 2 to 5 weeks of age. Macrophages were identified based on their expression of CD45⁺ SiglecF^−^F4/80⁺CD64^+^ (35).

We observed that the total count of CD64^+^F4/80^+^ macrophages per gram of muscle was significantly higher in dyW mice at all ages compared to WT mice, suggesting a robust and persistent inflammatory response in the dyW model (**Fig. 3A**). We further characterized the macrophage populations by measuring the frequencies of pro-inflammatory M1-like (Ly6C^+^CD206^−^) and anti-inflammatory M2-like (CD206^+^Ly6C^−^) macrophages. We observed a contrasting trend in the macrophage polarization between dyW and WT muscles (**Figs. 3B, 3C**). Most WT macrophages were pro-regenerative, anti-inflammatory (M2), with lower frequencies of M1. In contrast, dyW muscles display a distinct dynamic, with higher frequencies of M1 at the early stages of the disease (**Fig. 3B**), while the frequency of M2 remains constantly low over time (**Fig. 3C**). One significant pathway employed by inflammatory macrophages is inducible nitric oxide synthase (iNOS)-mediated production of nitric oxide (NO) from L-arginine (24). Macrophages that shift from M1 to an M2 phenotype typically undergo an increase in Arginase 1 expression that is concomitant with a reduction of iNOS expression (**Fig. 3D**). However, in dyW muscle the M2 macrophages displayed higher iNOS expression compared to Arg1, indicating impaired M1/M2 polarization and functional dysregulation of M2 macrophages (**Figs. 3 E-F**).

**Figure 3:**
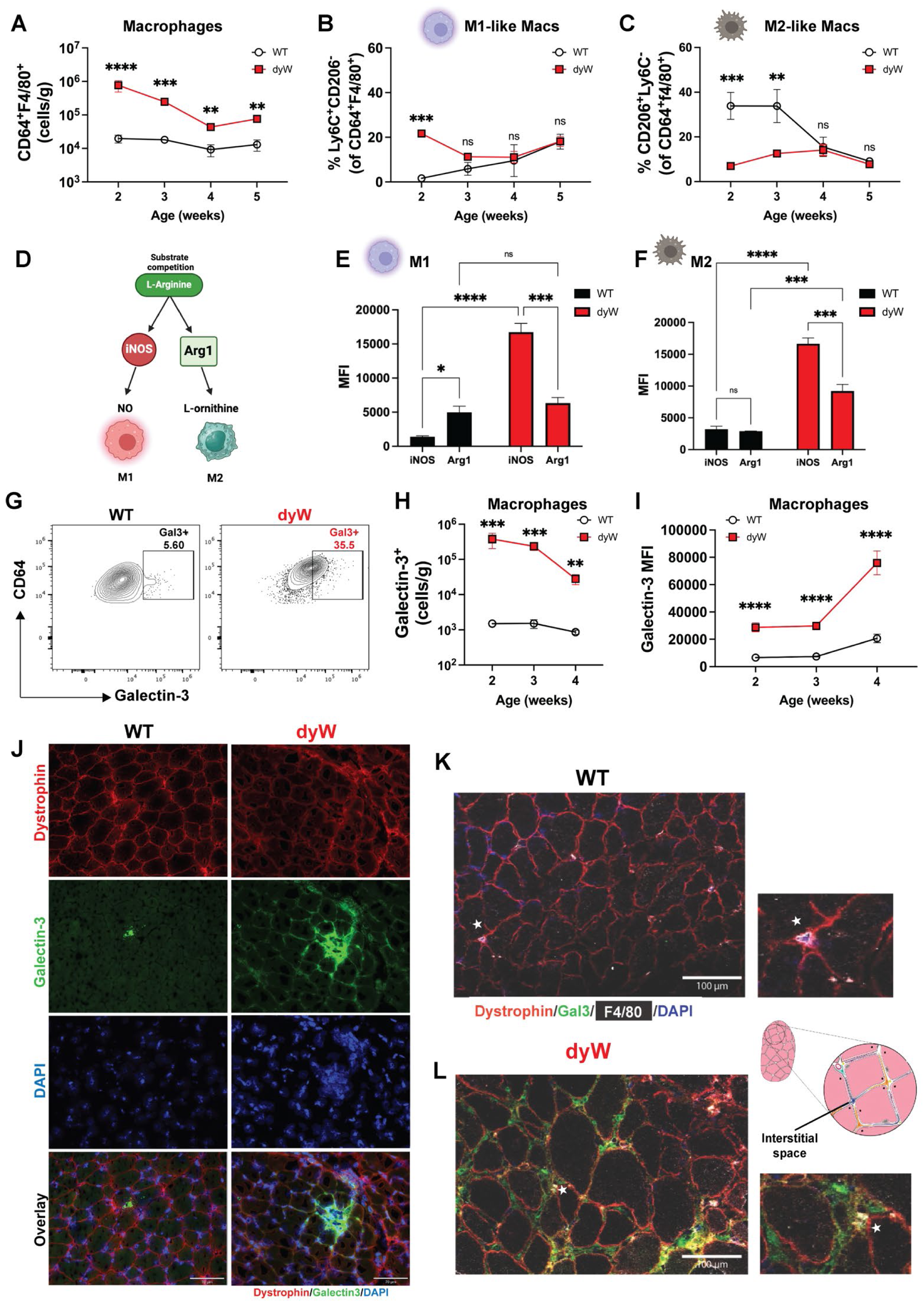
Characterization of Macrophages in dyW muscles. (A) Absolute cell counts of CD64^+^F4/80^+^ macrophages per gram of muscle at the indicated ages. (B) Frequency (%) of M1-like macrophages (Ly6C^+^CD206^−^ of CD64^+^F4/80^+^) as a percent of total macrophages at the indicated ages. (C) Frequency (%) of M2-like macrophages (CD206^+^Ly6C^−^ of CD64^+^F4/80^+^) as a percent of total macrophages at the indicated ages. (D) Brief overview of L-arginine metabolism in macrophage polarization. (E) Expression in mean fluorescence intensity (MFI) of iNOS and Arginase 1 (Arg1) in M1, and in (F) M2 macrophages from WT and dyW mice at 2 weeks of age. (G) Representative dot plots showing Galectin-3 expression in CD64^+^F4/80^+^ macrophages from WT and dyW mice. (H) Absolute cell counts of Galectin-3^+^ macrophages per gram of muscle at the indicated ages. (I) Expression in mean fluorescence intensity (MFI) of Galectin-3 in CD64^+^F4/80^+^ macrophages at the indicated ages. (J) Immunostaining against Galectin-3 (green), dystrophin (red), and nucleus (blue) in 2-weeks-old muscles (quadriceps). Galectin-3 is more pronounced in dyW muscles. Scale bar: 70 μm. (K-L) Immunostaining against Galectin-3 (green), Macrophage (F4/80 white), dystrophin (red), and nucleus (blue) in 2-weeks-old muscles (quadriceps) showing colocalization of galectin-3 with F4/80^+^ macrophage in dyW muscles. White asterisk indicates the colocalization and cartoon illustrates the interstitial space in the muscle. Scale bar: 100 μm. n= 5-7 mice/group. Data are presented as mean ± SEM. Statistical analysis was determined using two-way ANOVA with Tukey’s multiple comparisons. ns=not significant, *p ≤ 0.05, **p ≤ 0.01, *** p ≤ 0.005, **** p ≤ 0.001.

Because *Lgals3* (Galectin-3) was found to be one of the most upregulated genes in dyW muscles (log₂FC = 4.27, FDR p-value= 9.21×10^−88^; **Fig. 1C**) and a key upstream regulator in LAMA2-CMD (**Fig. 1F, Supplemental Table 3**), we examined its expression in macrophages. Galectin-3 is a β-galactoside-binding lectin highly expressed in immune cells, fibroblasts, and dystrophic muscle. It plays a crucial role in extracellular matrix remodeling, immune cell recruitment, and macrophage polarization (18, 27, 29, 31, 36–38). Given recent evidence linking Galectin-3⁺ macrophages to fibrosis in DMD mice (31), we investigated whether macrophages in LAMA2-CMD also express Galectin-3. Flow cytometry analysis revealed a higher frequency of Galectin-3⁺ macrophages in dyW muscles compared to WT controls (**Fig. 3G**). Quantification of Galectin-3⁺ macrophages per gram of tissue confirmed significantly elevated cell counts in dyW muscles throughout disease progression **(Fig. 3H**). Additionally, the expression of Galectin-3 by macrophages, measured by the mean fluorescence intensity (MFI), was significantly higher in dyW mice (**Fig. 3I**). Immunofluorescence staining showed elevated Galectin-3 expression in dystrophic dyW muscles (**Fig. 3J**), particularly in a subset of F4/80⁺ macrophages, while its expression was lower in WT muscle macrophages (**Fig. 3K**). These Galectin-3⁺ macrophages were predominantly localized within the interstitial space (**Fig. 3L**).

Collectively, these data suggest that an early surge in M1 macrophages drives the initial inflammatory response in LAMA2-CMD, while the reduced M2 macrophage frequencies indicate impaired resolution of inflammation. This imbalance likely contributes to sustained tissue damage and chronic inflammation, exacerbating disease progression. The increased presence of Galectin-3⁺ macrophages further support the role of persistent inflammation as a hallmark of LAMA2-CMD pathology. The distinct alterations in macrophage polarization and Galectin-3 expression between dyW and WT muscles underscore a dysregulated immune environment, highlighting macrophage modulation as a potential therapeutic strategy to mitigate disease progression.

### TD-139 treatment reduces galectin-3 expression on macrophages and reduces inflammatory markers in the dyW muscles

Galectin-3 is a key marker of chronic inflammation in macrophages and has been implicated in various inflammatory diseases (27–29, 36). Beyond its immune function, endogenous Galectin-3 plays a crucial role in muscle repair by regulating inflammation resolution and tissue regeneration (39). Given its dual involvement in immune regulation and muscle repair, we investigated the effects of Galectin-3 inhibition on the inflammatory profile and macrophage populations in LAMA2-CMD muscles. To do so, we used TD-139, a thio-digalactoside small-molecule inhibitor that targets the Galectin-3 carbohydrate-binding domain (**Fig. 4A**) and is currently being evaluated in clinical studies for its therapeutic potential in fibrotic, cancer and inflammatory disorders (26, 28–30, 40).

**Figure 4:**
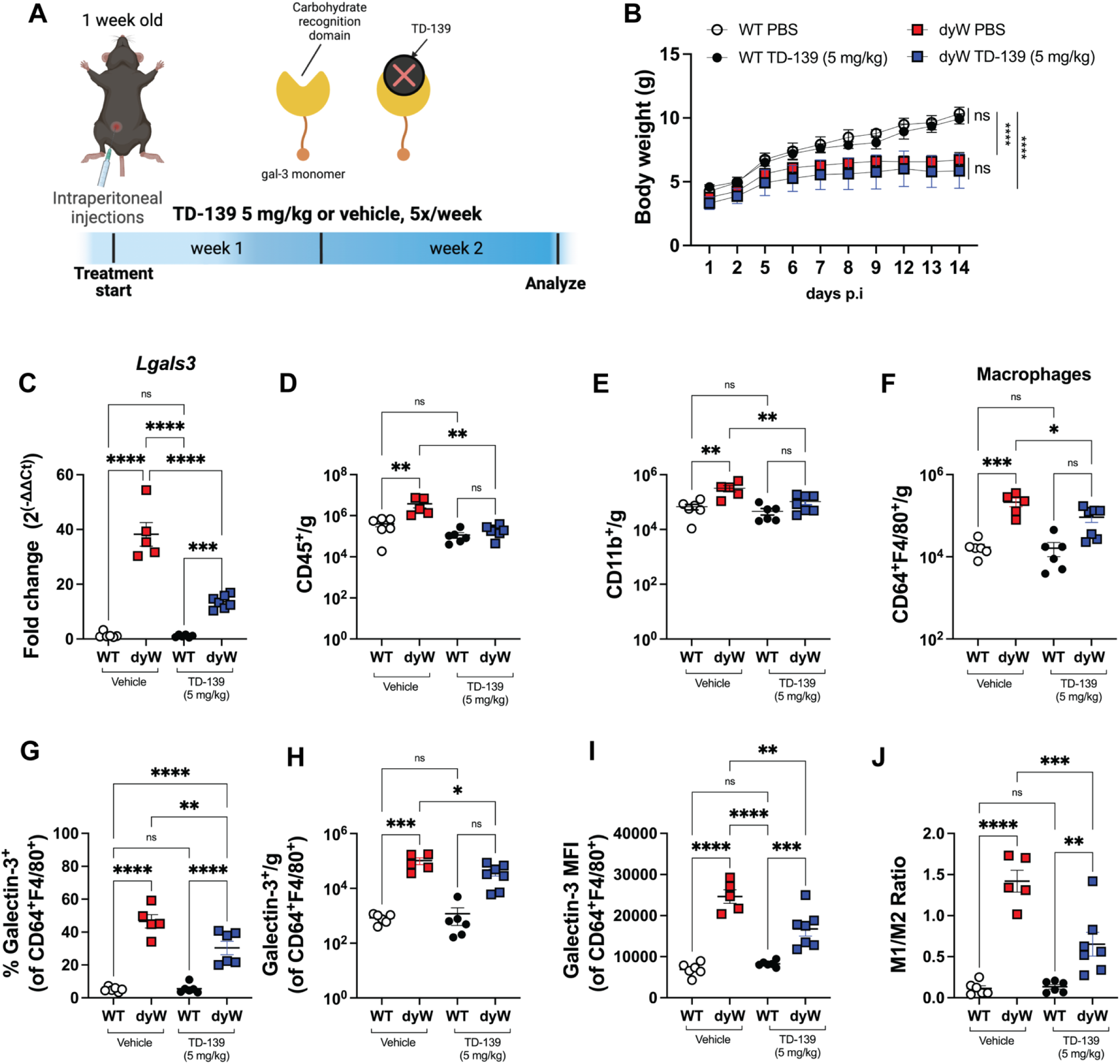
Impact of Galectin-3 inhibition on macrophages cells in dyW muscles. (A) Schematic representation of the experimental design. Mice received intraperitoneal (I.P.) injections of TD-139 at a dose of 5 mg/kg or vehicle (PBS+DMSO), administered five times per week for 2 weeks. (B) Body weight measurements in grams (g) over the 14-day period post-injections (p.i). (C) *Lgals3* mRNA levels in quadriceps. (D) Quantification of leukocyte (CD45^+^) cell counts per gram of tissue. (E) Absolute cell counts per gram of tissue of live myeloid cells (CD11b^+^). (F) Total macrophages (CD64^+^F4/80^+^ of live CD11b^+^ cells) per gram of tissue. (G) Frequency (%) and (H) absolute cell counts of live Galectin 3^+^ macrophages in the muscles. (I) Galectin 3 expression (MFI) in CD64^+^F4/80^+^ macrophages. (J) M1/M2 macrophage ratio. Data are presented as mean ± SEM. Results are pooled from two independent experiments. Each dot represents a single mouse. Statistical significance was determined using one-way ANOVA with Tukey’s multiple comparisons test (ns=non-significant, *p<0.05, **p<0.01, ***p<0.001, ****p<0.0001).

We treated the WT and dyW mice with intraperitoneal injections of TD-139 (5 mg/kg) or vehicle control (PBS) five times per week for 2 weeks (**Fig. 4A**). Throughout the duration of the treatment, we observed no significant differences in the body weight between animals receiving TD-139 or vehicle (**Fig. 4B**). Analysis of the quadriceps showed that TD-139 administration significantly reduced *Lgals3* mRNA levels (**Fig. 4C**). Subsequently, we quantified the number of lymphoid and myeloid cells using flow cytometry. As expected, the number of leukocytes (CD45^+^) per gram of tissue was significantly higher in the dyW mice compared to WT controls (**Fig. 4D**), indicating inflammatory infiltration. However, TD-139 treatment significantly reduced the number of CD45^+^ cells in dyW mice, without affecting the CD45^+^ cell number in WT mice (**Fig. 4D**). Importantly, we found that TD-139 treatment also significantly reduced the number of myeloid cells (CD11b^+^) in dyW muscles (**Fig. 4E**), specifically the population of macrophages (**Fig. 4F**). There was no change in the number of neutrophils or CD4^+^ and CD8^+^ T cells in dyW muscles (**Supplemental figures 3A-C**). Furthermore, the absolute cell counts and frequency (%) of Galectin-3^+^ macrophages were significantly reduced in dyW TD-139 treated mice compared to dyW vehicle controls (**Figs. 4G, H**). Among the total macrophages (CD64^+^F4/80^+^), Galectin-3^+^ macrophages accounted for an average of 5% in WT, but 45% in dyW (**Fig. 4G**). With TD-139 treatment, this frequency decreased to approximately 25% (**Fig. 4G**).

To determine the level of Galectin-3 protein expression on the macrophage cell surface, we measured the mean fluorescence intensity (MFI), which showed a significant decrease in dyW TD-139 treated mice, indicating effective reduction of Galectin-3 activity (**Fig. 4I**). Additionally, the M1/M2 macrophage ratio was significantly higher in dyW PBS-treated mice compared to WT controls, while TD-139 treatment significantly reduced the M1/M2 ratio in dyW mice, suggesting a shift towards a more anti-inflammatory macrophage profile (**Fig. 4J**).

Collectively, these results demonstrate that Galectin-3 inhibition with TD-139 significantly impacts the inflammatory profile and macrophage populations in LAMA2-CMD muscles. Specifically, TD-139 treatment reduces overall myeloid and macrophage infiltration, decreases the presence and activity of pro-inflammatory Galectin-3^+^ macrophages, and rebalances the M1/M2 macrophage ratio without affecting neutrophils or T cells. These findings warrant further evaluation to determine whether targeting Galectin-3 could serve as a therapeutic strategy to reduce chronic inflammation and improve disease outcomes in LAMA2-CMD.

### TD-139 modulates molecular pathways involved in muscle contraction, inflammation and fibrosis

Given the positive impacts of TD-139 treatment on the inflammatory milieu, we proceeded with assessing histopathology and transcriptomic profiles in the muscles of the treated animals. Hematoxylin and Eosin (H&E) staining on the dyW muscles isolated from vehicle-treated animals showed characteristic dystrophic pathology, including increased inflammatory cell infiltration and disorganized muscle fibers (**Supplemental Figure 4A**). In contrast, the muscles isolated from TD-139-treated dyW exhibited reduced inflammatory cell infiltration (**Supplemental Figure 4A**), which is consistent with the flow cytometry findings. Picrosirius Red staining revealed increased collagen deposition, indicative of fibrosis, in the dyW vehicle (**Supplemental Figure 4B**). However, there was no marked difference in collagen deposition between muscles isolated from vehicle- and TD-139-treated dyW (**Supplemental Figure 4B**). The short treatment duration (2 weeks) may not have been sufficient to drive detectable tissue remodeling or histological improvements.

To gain more insight into the molecular consequences of TD-139 treatment, we conducted bulk RNA sequencing analysis on quadriceps samples from WT and dyW mice, either treated or untreated with TD-139 at 3 weeks of age. The Venn diagram illustrates the differentially expressed genes (DEGs) across various groups comparing dyW and WT mice treated with either TD-139 or PBS (**Fig. 5A**). All DEGs can be found in **Supplemental Tables 4-5**.

**Figure 5:**
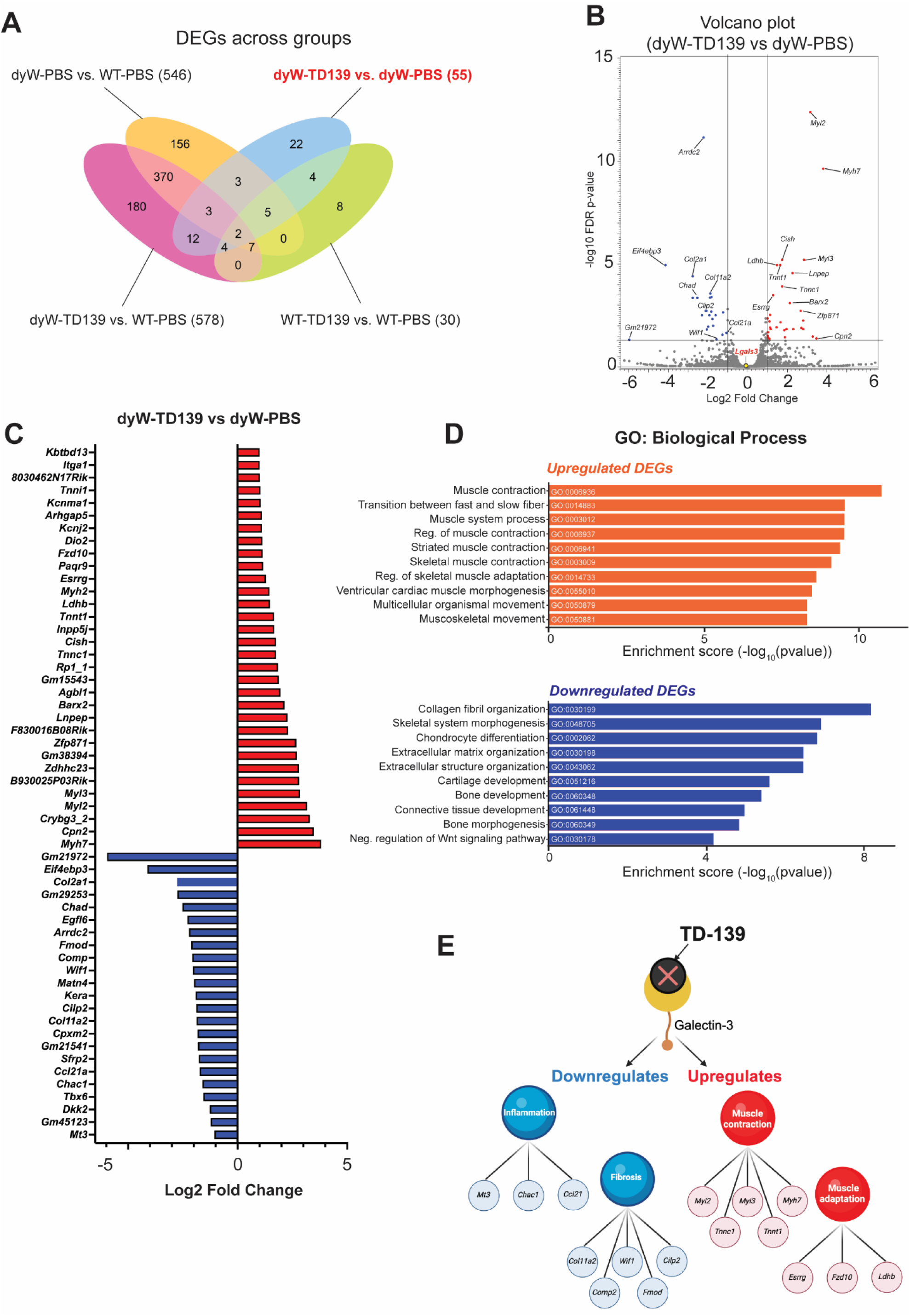
Impact of TD-139 on muscle transcriptome. (A) Venn diagram showing the differentially expressed genes (DEGs) at 3 weeks of age after TD-139 treatment. (B) Volcano plot displays DEGs in dyW-TD139 versus dyW-PBS with the most significant expressed genes highlighted. Red dots represent up-regulated genes and blue dots represent down-regulated genes, and the greys dots represents non-significant genes. The x-axis shows the log_2_ fold change, and the y-axis shows the −log10 FDR p-value. (C) Expression of the 55 genes differently expressed between dyW-TD139 vs dyW PBS. (D) Gene ontology (GO) analysis with a focus on biological processes displays the top 10 significantly enriched upregulated (orange) and downregulated (purple) processes in DEGs. (E) Summary of the processes affected by galectin-3 inhibition in dyW mice.

Subsequently, we sought to better understand the therapeutic effects of TD-139 in the context of the disease. Analysis of the DEGs between the dyW-TD-139 vs. dyW-PBS samples led to identification of 55 DEGs (Log_2_ FC ≥1, false discovery rate [FDR] ≤0.05; **Figs. 5B, 5C**). Within these, genes associated with muscle function/contraction (*Myh7*, *Myl2, Tnnc1*), adaptation (*Esrrg, Ldhb)*, and repair (*Myh7*, *Myl3, Cish*) were significantly upregulated upon TD-139 treatment (**Figs. 5B, 5C**). Gene ontology enrichment analysis (GO) also identified upregulated processes related to general muscle homeostasis and functions, including muscle contraction, transition between fast and slow fibers, and regulation of skeletal muscle adaptation (**Fig. 5D**). Collectively, these data indicate strengthened structural and functional components of skeletal muscle fibers as an impact of TD-139 treatment.

Conversely, TD-139 treatment downregulated genes associated with fibrosis, extracellular matrix remodeling, and inflammation, such as *Col2a1*, *Comp*, *Chad*, and *Ccl21a* (**Fig. 5B, 5C**). Indeed, the most significantly downregulated process on GO analysis (**Fig. 5D**) was collagen fibril organization, suggesting reduced extracellular matrix (ECM) deposition and remodeling, which contributes to decreased fibrosis. Processes such as ECM and extracellular structure organization further suggest TD-139’s ability to mitigate excessive ECM buildup and reduce tissue fibrosis. Additionally, the inhibition of processes such as skeletal system morphogenesis, chondrocyte differentiation, and negative regulation of Wnt signaling pathway suggests that TD-139 mitigates maladaptive tissue remodeling.

Collectively, our data suggest that inhibition of Galectin-3 via TD-139 treatment may improve the dystrophic muscle environment by regulating pathways associated with muscle contraction, adaptation, inflammation, and fibrosis (**Fig. 5E**).

## Discussion

The motivation of this study was two-fold: first, prior knowledge pertaining to resident immune cells in LAMA2-CMD muscles have relied heavily on qualitative or semi-quantitative methods, such as histopathology and immunostaining. Moreover, despite its clear presence in muscle biopsies from LAMA2-CMD-affected individuals and mouse models, inflammation remains an underexplored area in the development of LAMA2-CMD therapies. Therefore, we sought to investigate the molecular players in the LAMA2-CMD muscle inflammatory milieu by integrating unbiased transcriptomic profiling with comprehensive flow cytometry-based immunophenotyping and subsequently use the knowledge to inform pharmacological intervention.

We laid the foundation for this effort by analyzing the transcriptomic profiles of muscles isolated from dyW mice at 5 weeks of age. Strikingly, genes involved in leukocyte processes including migration, activation, proliferation, cell-cell adhesion, and chemotaxis were significantly upregulated. Chief among these upregulated genes was the *Lgals3* gene encoding for Galectin-3, a β-galactoside-binding lectin expressed in various tissues, including muscles (39, 41) and immune cells, such as macrophages, where it plays a role in both acute and chronic inflammation (31). Importantly, such inflammation-heavy processes were absent in the transcriptomic profiles of dyW muscles at E17.5 (42), marking a distinction between disease onset vs. advanced stages of the disease. Nevertheless, we also detected changes in developmental myosin heavy chain (MyH) and -light chain (Myl) isoforms, such as *Myh3, Myl4,* and *Myh8*, as well as *Spp1* (Osteopontin) and *Sln* (Sarcolipin). This observation is similar to previously reported transcriptomic findings in the aforementioned embryonic dyW (42) and 8-week-old dy2j (43) mice, as well as proteomic analysis in muscles isolated from 4-week-old dy3k mice (44), indicating that impaired muscle regeneration and fibrosis are consistent across different disease mouse models and ages.

Alongside the upregulated inflammation-related processes, we also observed downregulation of processes that may impair muscle contractility and functions, such as ion and potassium transmembrane transport, muscle system process, regulation of relaxation in muscle, in the transcriptomic profile of 5-week-old dyW quadriceps. Dysfunction of the ion homeostasis and potassium transmembrane transport pathway has been previously reported in the brain of dyH mice, suggesting a broader systemic impairment of these pathways in LAMA2-CMD (45). Energy reserve-, glycogen- and glucan metabolic processes were also downregulated in dyW muscles, indicating a potential impact on muscle endurance. Such findings are in accordance with a previous proteomic analysis in dy3K muscles, which showed reduced expression of proteins primarily involved in metabolic pathways such as glycolysis, fatty acid oxidation, and oxidative phosphorylation (44).

Building on the enriched inflammation-related biological processes identified on the transcriptomic profiling, we further dissected the population of immune cells in the muscles. We employed flow cytometry-based immunophenotyping, which allows simultaneous detection and quantification of multiple immune cell properties including their identity, activation state, and subsets, within a single sample, thus bypassing limitations in the number of markers analyzed with qualitative or semi-quantitative methods such as histopathology or immunostaining (8, 15, 17, 18, 46). Our immunophenotyping analysis revealed that dyW muscles exhibit an early transient yet robust leukocyte (CD45^+^) infiltration, which eventually declined by 5 weeks, reflecting the notion that the inflammatory response is triggered by muscle damage early during disease onset in LAMA2-CMD (8, 15, 17). In addition, macrophages, here defined as CD45⁺F4/80⁺SiglecF^−^CD64^+^, were found to be the most predominant myeloid cell type in the dystrophic muscle. Previous studies have relied on CD11b or CD68 markers to identify clusters of macrophages on muscle sections using immunostaining (8, 15, 17). However, these markers are not sufficient to characterize macrophages, because CD11b is expressed by other immune cell types, including neutrophils, monocytes, eosinophils, natural killer cells, and dendritic cells and these cell types are also present in dystrophic muscles (47).

In addition to better defining the macrophage population, we were also able to detect impaired M1/M2 polarization in the dyW quadriceps. M2 macrophages, essential for muscle repair, remained low and unchanged over time, indicating macrophage imbalance in LAMA2-CMD. While M1 macrophage numbers did not increase over time, they showed elevated iNOS expression, suggesting persistent activation which contributes to inflammation. This dynamic pattern is similar to that observed in Duchenne muscular dystrophy (DMD), where M1 macrophages drive acute inflammation and tissue damage, while an insufficient M2 macrophages frequency hinders the resolution of inflammation and muscle repair (9, 24, 25).

The integration of rich transcriptomics- and highly quantitative immunophenotyping data revealed two key findings: a significantly high expression of *Lgals3* and its position as a main upstream regulator in the transcriptomic signatures, as well as impaired macrophage polarization. Further analysis with immunostaining and flow cytometry showed that macrophages in LAMA2-CMD muscles are enriched for Galectin-3, both in terms of expression and localization. High *Lgals3* expression has been reported in the muscles of both dyW and dy3k mice (18, 32); however, the direct spatiotemporal connection with macrophages in LAMA2-CMD has not been made prior to our study.

Galectin-3 contributes to the initiation and amplification of the acute inflammatory response by recruiting macrophages to injury sites and perpetuates a state of chronic inflammation through activation of proinflammatory pathways. Several pharmacological inhibitors of Galectin-3 that block its carbohydrate recognition domains exist, including TD-139, which is currently being investigated in clinical studies for its therapeutic potential in fibrotic, cancer and inflammatory disorders (26, 28, 29, 40). In this study, we showed that systemic TD-139 administration effectively reduced overall myeloid and macrophage infiltrations, decreased the presence and activity of pro-inflammatory Galectin-3^+^ macrophages, and rebalanced the M1/M2 macrophage ratio in the dyW quadriceps. It is important to acknowledge that the experiment we conducted here was not designed to be an extensive, long-term pre-clinical study. Nonetheless, the transcriptomic analysis on TD-139-treated dyW muscles showed upregulation of genes involved in muscle contraction and adaptation and downregulation of those involved in ECM organization. Given that TD-139 treatment did not alter neutrophil or T-cell populations in either dyW or WT muscles, these data suggest that its effects are largely macrophage-dependent. Thus, by inhibiting Galectin-3, TD-139 supports a pro-regenerative macrophage profile, which may eventually improve muscle homeostasis in LAMA2-CMD.

While Galectin-3 is not a new player in fibrotic and inflammatory diseases, per se, it is notoriously known to be a ‘hard-to-tame’ target (48). Transgenic mice lacking both LAMA2 and Galectin-3 (dy3K/GAL double knockout) showed no significant improvements in pathology, muscle function, or inflammation (18). In contrast, pharmacological inhibition of Galectin-3 with TD-139 pointed towards reduced inflammation in dyW mice. We reason that, unlike genetic ablation that leads to the complete loss of Galectin-3 from the embryonic stage and may activate compensatory mechanisms, TD-139 provides partial and temporally controlled inhibition, allowing preservation of Galectin-3’s beneficial roles in tissue repair and resolution of acute inflammation (39). Our findings suggest that TD-139 not only reduces pathological inflammation but also activates transcriptional programs that support muscle regeneration and function, while still preserving the beneficial roles of endogenous Galectin-3 in the muscle. Although histological analysis did not show significant reduction in collagen deposition, this likely reflects the short (e.g., two-weeks) treatment duration, as tissue remodeling typically requires extended periods to become histologically evident. Future studies would focus on investigating longer treatment regimens, dosages, and assess functional outcomes, such as muscle strength and endurance, to provide a more comprehensive evaluation of therapeutic efficacy. Additionally, combining Galectin-3 inhibition with a variety of genetic therapies in development for LAMA2-CMD, including but not limited to upregulation of LAMA1 (49–52), linker proteins (53–55), or micro-laminin (56), may yield synergistic benefits.

While Galectin-3 and macrophages largely became the focus of our study, several additional findings are worth emphasizing. Our immunophenotyping effort showed an increase in T-regulatory cells (CD3^+^CD4^+^Foxp3^+^; Tregs) at early stages of the disease, *i.e.,* at 2 and 3 weeks, followed by a subsequent decline. These Tregs exhibited an activated phenotype, as evidenced by the elevated frequencies of GITR and KLRG1 marker positivity in Tregs from dyW muscle compared to WT muscle. Interestingly, Tregs are also elevated in DMD (11). Studies in dystrophin-deficient mdx mice have shown that depletion of Tregs exacerbates muscle inflammation and injury, which was characterized by an enhanced interferon-gamma (IFN-γ) response and activation of pro-inflammatory (M1) macrophages (11). Given the early increase in Tregs in dyW mice, we speculate that signals arising from muscle injury initially drive the expansion of Tregs. However, in the context of chronic inflammation, Tregs might fail to regulate exacerbated inflammation in dyW muscle. Future work focusing on the mechanism underlying Treg dysfunction in LAMA2-CMD could provide valuable insights for therapeutic interventions. To our knowledge, this study is the first to employ immunophenotyping by flow cytometry as the primary method for characterizing the heterogeneity of immune cell populations in LAMA2-CMD muscles, thereby providing an additional layer of information on what transpires in the dystrophic muscle environment.

Overall, our study and others emphasize that LAMA2-CMD pathology is complex, ranging from metabolic alterations to inflammation. This complexity should be taken into consideration when designing and analyzing therapeutic interventions. With regards to inflammation in LAMA2-CMD, the question remains: what is the best way to achieve a balance where the detrimental effects of inflammation are reduced, and the beneficial aspects are preserved? Immunodeficient dyW mice lacking the adaptive immune response, *i.e.*, CD3 and B cells, exhibit impaired muscle regeneration (57). Connolly et al observed improvements when treating LAMA2-CMD mice with prednisolone (58), a medication that suppresses general inflammation. However, it is known that long-term use can be detrimental due to significant side effects in humans (59). Moreover, the complete lack of a key player like Galectin-3 in LAMA2-CMD may not be therapeutically advantageous (18), despite evidence that elevated Galectin-3 is strongly associated with detrimental inflammation, including but not limited to our study presented here. Pharmacological inhibition with TD-139 effectively reduces inflammation, modulates macrophage polarization, and enhances muscle contraction and adaptation transcriptomic profiles, positioning Galectin-3 as a viable therapeutic target.

Moving forward, we propose that Galectin-3 should not be considered as a standalone player. As a lectin, Galectin-3 exerts its action by modulating the biological activity of its binding partners. In this context, investigating the presence and expression of Galectin-3 and its interactants in LAMA2-CMD muscles, both in patient and animal models, is essential. This would enable the construction of a comprehensive map of the Galectin-3 interactome involved in disease processes – an approach that could inform the development of more targeted and effective pharmacological strategies for inflammation and immune dysregulation in diseases, including but not limited to LAMA2-CMD.

## Methods

### Sex as a biological variable

Sex as a biological variable was considered by making use of both male and female animals. Findings were similar for both sexes.

### Mice

dyW mice (B6.129S1(Cg)-*Lama2^tm1Eeng^*/J; strain #013786) were purchased from the Jackson laboratory, housed and bred in the University of Pittsburgh’s Division of Laboratory Animal Research facility at the Rangos Research Building, UPMC Children’s Hospital of Pittsburgh under specific pathogen-free conditions.

### TD-139 treatment

Mice were treated with TD-139 (Selleckchem, S0471) at a dose of 5 mg/kg via intraperitoneal injection, 5 times per week for a period of 2 weeks. Control mice were administered PBS + 0.05% DMSO for the same frequency and duration.

### Cell isolation and flow cytometry

The left and right hindlimb muscles from each individual mouse were pooled, preserved in DMEM (Gibco), and kept on ice until tissue processing. Single cells were then isolated using the Skeletal Muscle Dissociation Kit (Miltenyi Biotec, Bergisch Gladbach, Germany) and the gentleMACS Octo Dissociator with heaters (Miltenyi Biotec), following manufacturer’s instructions. Briefly, under sterile conditions, the muscle tissue was minced into small pieces and transferred to a gentleMacs C tube containing 2.5 mL of a mix of enzymes. The tubes were then placed into the gentleMacs Octo Dissociator with heaters and processed using the 37C_mr_SMDK_1 program, incubating the samples at 37 °C for 1 hour under continuous rotation. Following incubation, the samples were centrifugated at 300 g for 10 min at 4°C. The cell pellet was resuspended in 10 mL of DMEM (Gibco) and filtered through a 70 μm cell strainer. Cells were counted using a hemocytometer, and yields were reported as the total number of viable cells. Subsequently, cells were then aliquoted at 2×10^6^ cells/well in a 96-well plate and immediately stained with Live-Dead Fixable blue (BUV496; Thermo) at 1:1000 for 15 minutes at room temperature, protected from the light. Non-specific antibody binding was blocked by TruStain FcX™ CD16/CD32 (Clone 93, Biolegend). The cells were stained with a surface antibody cocktail containing: Anti-CD4 (BUV395, clone GK1.5, BD Biosciences), GITR (BUV563, clone DTA-1, BD Biosciences), Ly6C (BUV661, clone HK.1.4.rMab, BD Biosciences), KLRG1 (BUV737, clone 2F1, BD Biosciences), Ly6G (BUV737, clone 1A8, BD Biosciences), CD8 (BUV805, clone 53-6.7, BD Biosciences), CD68 (BV421, clone FA-11, Biolegend), CD11b (BV570, clone M1/70, Biolegend), CD206 (BV605, clone C068C2, Biolegend), CD64 (FITC, clone X54-5/7.1, Biolegend), SiglecF (PE-Dazzle-594, clone S17007L, Biolegend), Galectin-3 (PE-Cy7, clone M3/38, Biolegend), CD3 (PE-Cy5, clone 17A2, Biolegend), CD45 (APC-Cy7, clone 30-F11, Biolegend), for 30 minutes at 4°C, fixed for 20 minutes with 1% paraformaldehyde, and permeabilized with True-Nuclear Transcription Factor (Biolegend; Cat No. 424401) for 45 min at room temperature. Cells were stained intracellularly with an antibody cocktail containing Foxp3 (PE, clone MF-14, Biolegend), iNOS (APC, clone W16030C, Biolegend), Arginase 1 (PE-Cy7, Clone A1exF5, ThermoFisher) for 20 min at room temperature, washed, and analyzed immediately. Flow cytometry was performed using a Cytek Aurora machine (Cytek Biosciences, Rangos Flow Cytometry Core Laboratory). Flow Cytometry Standard (FCS) files were analyzed using FlowJo software for Macintosh (version 10.1). Gating strategies are shown in **Supplemental Figures 1A-C**.

### Histology, immunostaining, and imaging

Muscles were isolated, preserved in liquid nitrogen-chilled isopentane, and stored at −80 °C until further use. Frozen muscles were embedded in OCT and sectioned at 8 μm thickness using a cryostat at −20 °C. Sectioned tissues were mounted on slides and stained using Hematoxylin and Eosin (H&E) and Picrosirius red. For immunostaining, frozen muscle sections were fixed in ice-cold methanol for 10 min, washed in PBS and blocked for 1 h in 5% goat serum in PBS. Primary antibodies were incubated overnight at 4°C, while secondary antibodies were incubated for one hour at room temperature. The following primary and secondary antibodies were used: mouse anti-dystrophin (1:100, DHSB-MANDRA11), rabbit anti-galectin-3 (1:100, Cell Signaling #E7B6R), rat anti-F4/80 (1:50, Abcam, ab6640), goat anti-mouse Alexa-594 (1:1000, Thermofisher, A11005), goat anti-rabbit Alexa-488 (1:1000, Thermofisher, A11008), goat anti-rat Cy5 (1:500, Thermofisher, A10525).

Imaging of muscle tissue sections was performed using an ECHO Revolve microscope or a Leica Stellaris 5 confocal laser scanning microscope equipped with 20x or 63x oil immersion objectives using LASX software. Final figures were composed using ImageJ.

### RNA isolation and quantitative polymerase chain reaction (qPCR)

Total RNA was extracted from mouse tissues using TRIzol reagent (Thermo) following the manufacturer’s protocol. Using 1 µg of total RNA, cDNA was synthesized using the iScript Reverse Transcriptase Supermix kit (BioRad, Cat# 1708841) as per the manufacturer’s guidelines. 2 µl of 1:5 diluted cDNA was used for the Quantitative PCR (qPCR) carried out using the 2X SYBR Green Fast qPCR Mix kit (ABclonal, Cat# RM21203) on a C1000 Touch Thermal Cycler (BioRad, USA). The following primers were used: *Lgals3*_Fwd 5’-aacacgaagcaggacaataactgg-3’, *Lgals3*_Rev 5’-gcagtaggtgagcatcgttgac-3’, *Gapdh*_Fwd 5’-acatggcctccaaggagtaagaa-3’ *Gapdh*_Rev 5’-gggatagggcctctcttgct-3’. *Gapdh* expression was used as an internal control to normalize the relative expression levels of the genes of interest. The 2^ΔΔCt^ method was applied to determine the relative fold-change in gene expression.

### RNA sequencing and pathway enrichment analysis

Total RNA was isolated from mouse tissues using TRIzol reagent (Thermo), according to the manufacturer’s instructions. After extraction, the samples were submitted to the Health Sciences Sequencing Core at the UPMC Children’s Hospital of Pittsburgh for further processing. RNA quality was assessed using the Agilent Bioanalyzer 5300 Fragment Analyzer (Agilent Technologies). cDNA libraries were prepared using the Illumina Stranded mRNA library preparation kit (Illumina), and sequencing was performed on the NextSeq 2000 platform with pair-end 58 bp reads. Analysis of sequence reads, including quality control, mapping and generation of tables of DEGs, heatmaps and volcano plots were performed using QIAGEN licensed CLC Genomic Workbench software (version 22.0.1). Gene ontology analysis was performed using the SRplot database (33). For RNA-seq analysis, genes were selected using a cutoff of Log_2_ FC ≥1.5 and FDR p-value ≤0.05 in **Fig. 1**, and Log_2_ FC ≥1 and FDR p-value ≤0.05 in **Fig. 5**. Upstream regulator analysis and Top 10 diseases annotations and functions were carried out using QIAGEN-licensed Ingenuity Pathway Analysis (IPA) software.

### Statistical analyses

Shapiro-Wilk normality test was performed to check for normal distribution of the data. The statistical significance between groups was obtained with one-way or two-way ANOVA with Tukey’s post hoc correction test to compare more than two sets of normally distributed data. Statistical significance was defined as P value <0.05. Statistical analyses were performed using GraphPad Prism software Version 10.2.3.

### Study approval

Animal protocols and procedures were approved by the University of Pittsburgh Institutional Animal Care and Use Committee (IACUC) under protocol number 22061237.

## Data availability

Raw data of RNA-Sequencing will be available in the NCBI’s Gene Expression Omnibus database (GEO) after April 1^st^, 2025 release date.

## Author contributions

YKTdM and DUK designed the study; YKTdM, JL, JQC-Z, MAJ, RNR conducted experiments; YKTdM, JL, JQC-Z acquired and analyzed data; YKTdM wrote the manuscript with critical input from DUK and all the coauthors.

## Acknowledgements

Members of the Kemaladewi Lab are acknowledged for the critical feedback and technical assistance on this project and manuscript. We thank Dr. Sameer Agnihotri (UPMC Children’s Hospital of Pittsburgh), Dr. Krishna Prasadan (Imaging Core, Dept. of Pediatrics, Univ. of Pittsburgh School of Medicine), Dr. Abbe Vallejo, Dr. Radha Gopal, and Josh Michael (Flow Cytometry Core, Dept. of Pediatrics, Univ. of Pittsburgh School of Medicine), Dr. William MacDonald (Health Sciences Sequencing Core, Univ. of Pittsburgh School of Medicine) for their help with tissue lysis, confocal imaging, flow cytometry, and RNA sequencing, respectively. The CLC Genomics Workbench and Ingenuity Pathway Analysis software, licensed through the Molecular Biology Information Service of the Health Sciences Library System at the University of Pittsburgh, were used for data analysis. This work is supported by a postdoctoral trainee grant from the Research Advisory Committee, UPMC Children’s Hospital of Pittsburgh (to YKTdM), Development Fund from Dept. of Pediatrics, Univ. of Pittsburgh School of Medicine, AFM-Telethon (23136) and Cure CMD Research Grants, NIH R01 AR078872, and NIH Director’s New Innovator Award DP2 AR081047 (to DUK).

---

**Supplemental Figure 1:**
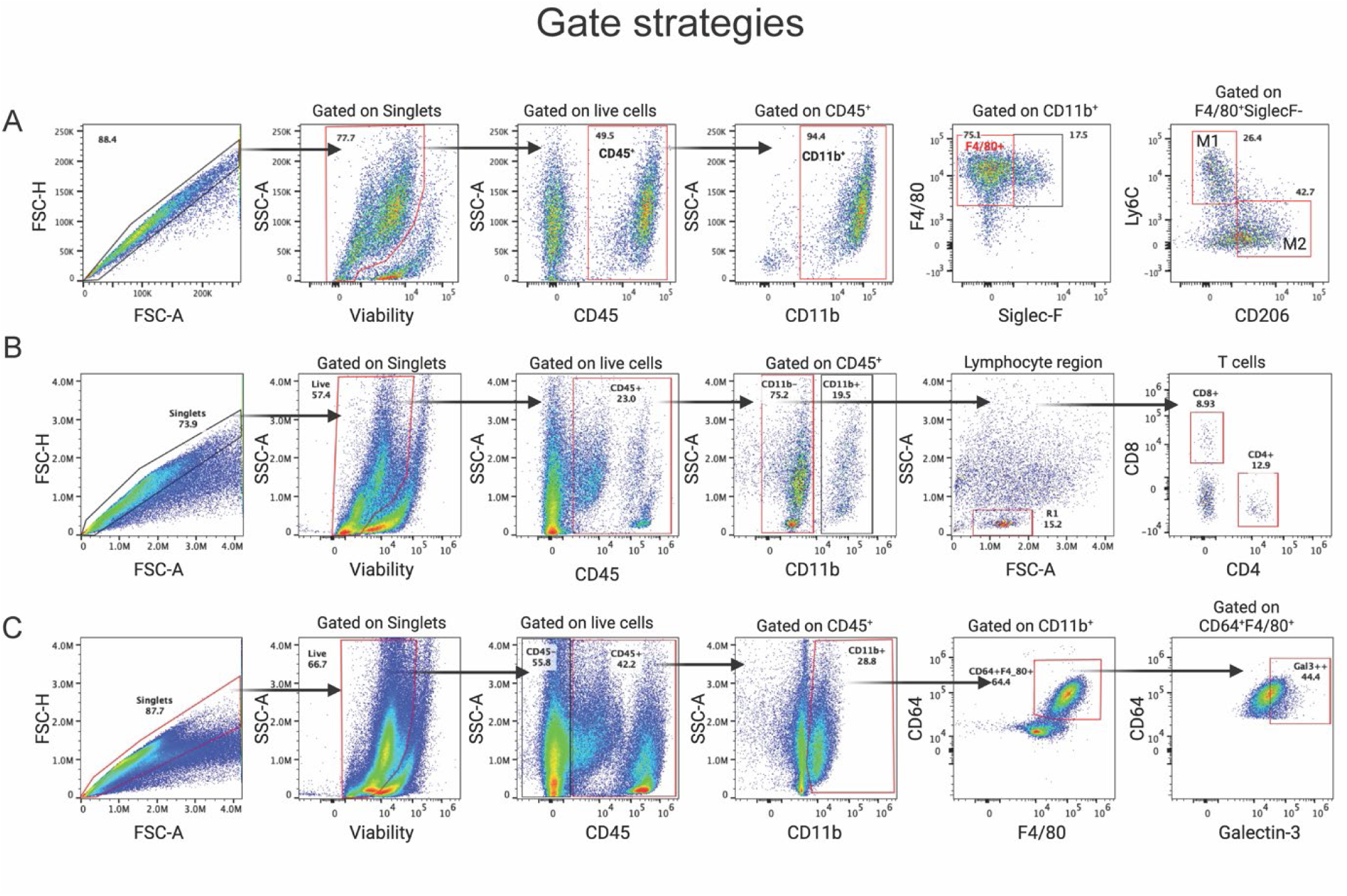
Gating strategies used to identify targeted populations in the muscles. (A) Gating strategy for Macrophages. Single cell suspensions were gated based on their scatter to identify singlets followed by exclusion of dead cells. Leukocytes were identified based on CD45 staining. Myeloid cells were identified by CD45^+^CD11b^+^. Macrophages were identified by F4/80^+^ and SiglecF^−^ staining. M1 macrophages were identified as F4/80^+^SiglecF^−^Ly6C^+^CD206^−^ and M2 macrophages as F4/80^+^SiglecF^−^CD206^+^Ly6c^−^. (B) CD4 and CD8 T cells were identified by gating on CD11b^−^ (negative) cells. (C) Galectin-3+ Macrophages were identified by the expression of CD11b^+^CD64^+^F4/80^+^Galectin3^+^ markers.

**Supplemental Figure 2:**
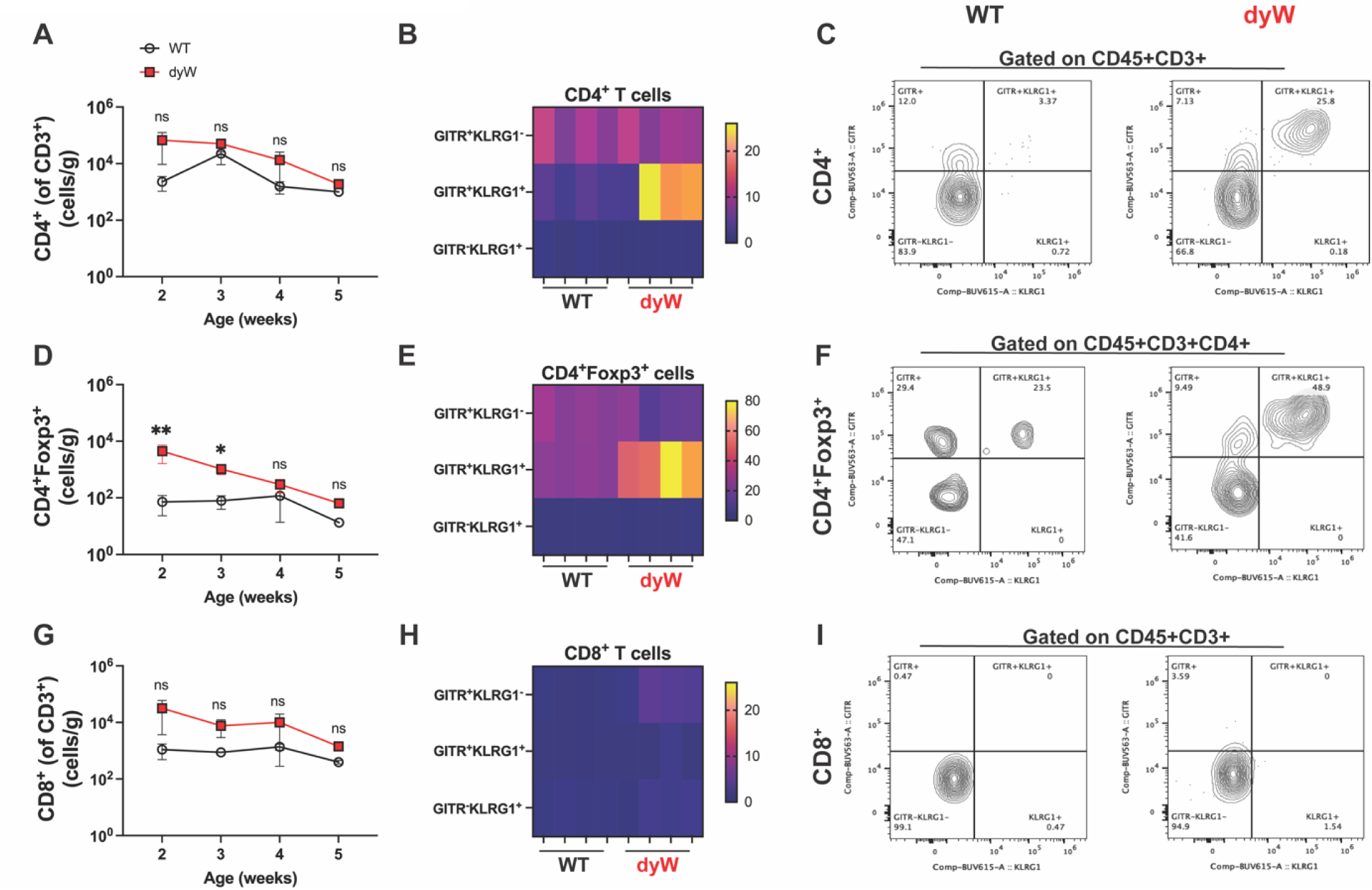
Characterization of CD3^+^ T lymphocytes in dyW muscles. (A) Absolute CD4+CD3+ cell counts in WT and dyW mice. (B) Expression of activation markers GITR and/or KLRG1 at 2 weeks in CD4+ cells. (C) Representative dot plots showing increased frequency of CD4+ T cells expressing GITR+KLRG1 at 2 weeks. (D) Absolute Tregs CD4+Foxp3+ cell counts in WT and dyW mice. (E) Expression of activation markers GITR and/or KLRG1 at 2 weeks in Tregs. (F) Representative dot plots showing increased frequency of CD4+Foxp3+ cells expressing GITR+KLRG1 at 2 weeks. (G) Absolute Tregs CD8+CD3+ cell counts in WT and dyW mice. (H) Expression of activation markers GITR and/or KLRG1 at 2 weeks in CD8+ cells. (I) Representative dot plots showing the frequency of CD8+ cells expressing GITR and/or KLRG1 at 2 weeks. n= 5-7 mice/group. Data are presented as mean ± SEM. Statistical significance was determined by one-way ANOVA with Tukey’s multiple comparisons test (ns= non-significant, *p<0.05, **p<0.01).

**Supplemental Figure 3:**
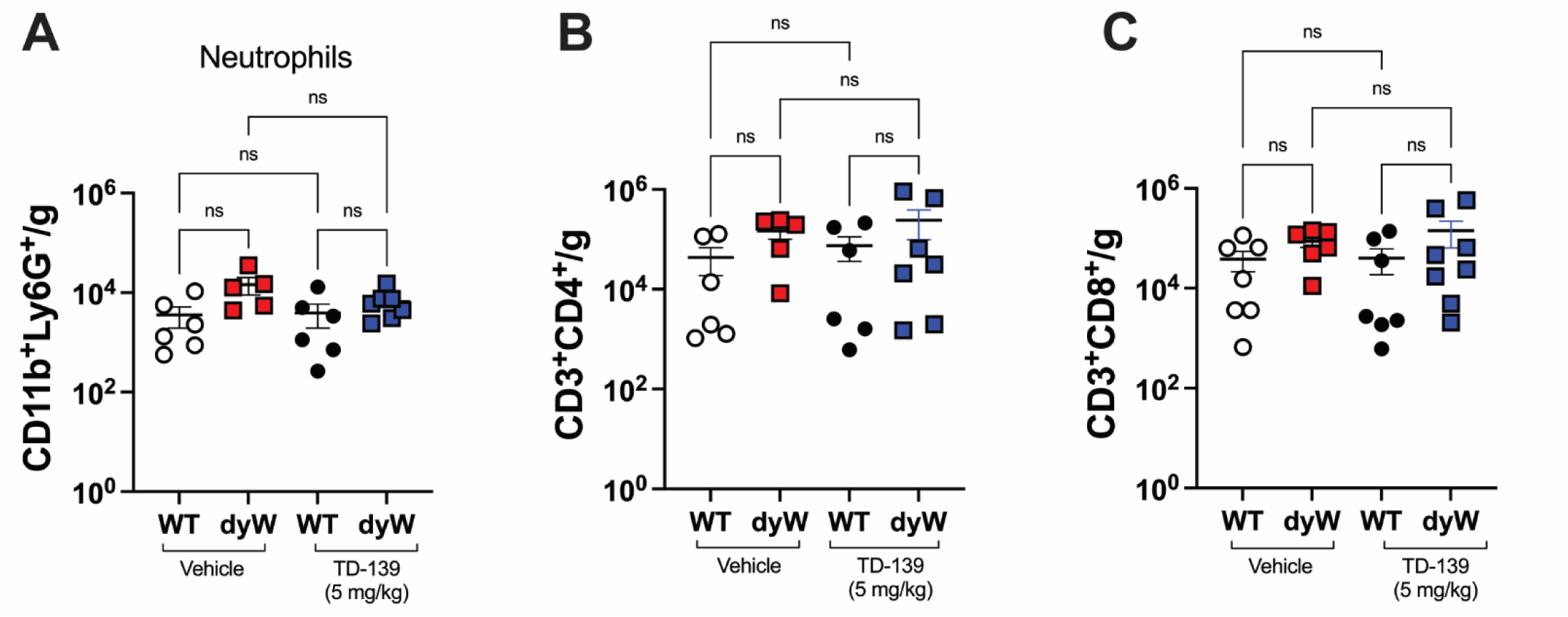
TD-139 does not alter Neutrophils and T cells counts in the muscles. (A) Absolute live Neutrophils (CD11b^+^Ly6G^+^) counts in quadriceps from WT and dyW treated either with vehicle or TD-139. (B) Absolute live CD4^+^ T cells and (C) CD8^+^ T cells counts in quadriceps from WT and dyW treated either with vehicle or TD-139. n= 5-7 mice/group. The results are pooled from two independent experiments. Data are presented as mean ± SEM. Statistical significance was determined by one-way ANOVA with Tukey’s multiple comparisons test (ns= non-significant).

**Supplemental Figure 4:**
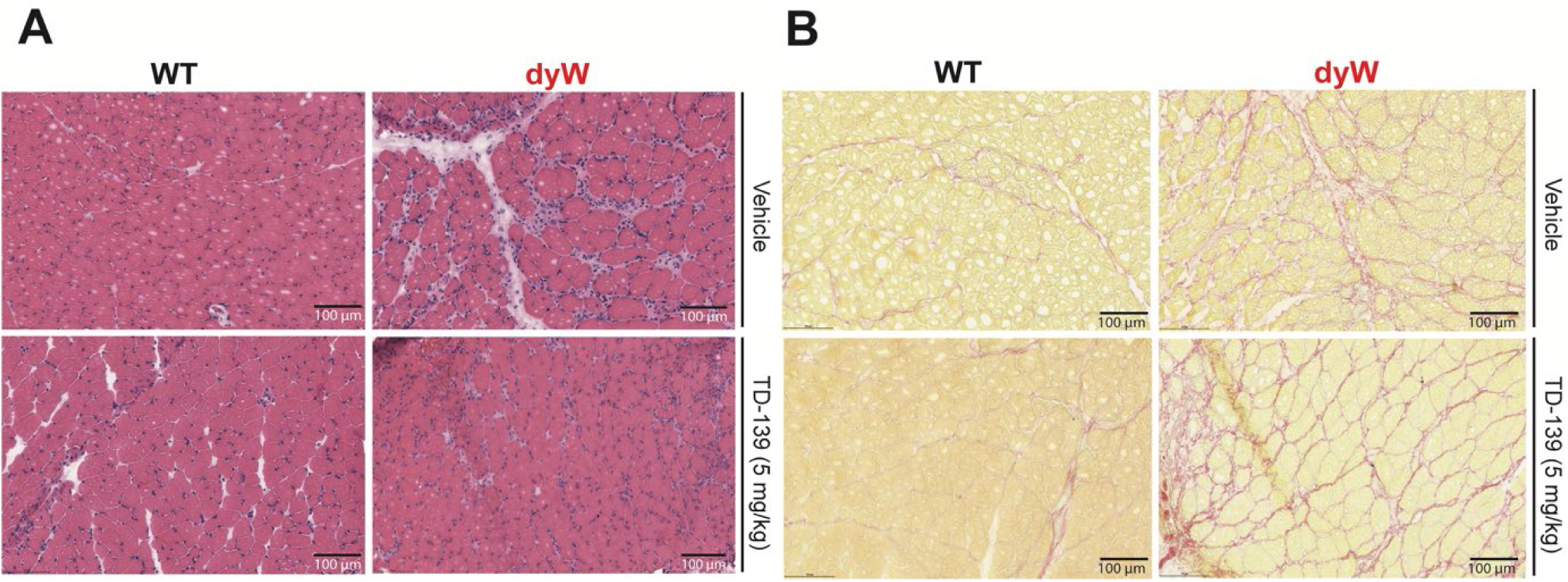
Impact of TD-139 on muscle histopathology. **(A)** Representative H&E and **(B)** Picrosirius red staining images of quadriceps muscle frozen section from WT, dyW vehicle and TD-139 treated (scale bar 100 μm).

